# Allele-specific transcription factor binding as a benchmark for assessing variant impact predictors

**DOI:** 10.1101/253427

**Authors:** Omar Wagih, Daniele Merico, Andrew Delong, Brendan J Frey

## Abstract

Genetic variation has long been known to alter transcription factor binding sites, resulting in sometimes major phenotypic consequences. While the performance for current binding site predictors is well characterised, little is known on how these models perform at predicting impact of variants. We collected and curated over 132,000 potential allele-specific binding (ASB) ChIP-seq variants across 101 transcription factors (TFs). We then assessed the accuracy of TF binding models from five different methods on these high-confidence measurements, finding that deep learning methods were best performing yet still have room for improvement. Importantly, machine learning methods were consistently better than the venerable position weight matrix (PWM). Finally, predictions for certain TFs were consistently poor, and our investigation supports efforts to use features beyond sequence, such as methylation, DNA shape, and post-translational modifications. We submit that ASB data is a valuable benchmark for variant impact on TF binding.

## Introduction

One of the primary mechanisms contributing to the regulation of gene expression is the binding of transcription factors (TFs) to regulatory genomic elements. Differential gene expression can drive and contribute to almost every aspect of disease phenotypes. Understanding the intricate process of TF-DNA binding can, therefore, provide mechanistic hypotheses for variants and propel the discovery of novel therapies. Fortunately, through the aid of high throughput techniques such as chromatin immunoprecipitation followed by sequence (ChIP-seq), systematic evolution of ligands by exponential enrichment (SELEX) and protein binding microarrays (PBMs), the binding specificities of many TFs have been exhaustively catalogued over the past decade^1^.

Genetic variation falling within specificity determinants of transcription factor binding sites (TFBSs) can alter binding by introducing novel binding sites or diminishing existing binding sites, often resulting in a substantial impact on molecular phenotypes through changes in gene expression. Although experimental approaches have utilised ChIP-seq to map variants to molecular-level traits such as TF-binding^2,3^, these approaches are costly and cannot yet be routinely applied to the sizeable quantity of genetic variation data available. As such, much effort has gone into modelling TF-DNA binding *in silico*. The current standard for predicting TFBSs is the position weight matrix, owing to its simplicity and intuitiveness^4^. More recent machine learning, and specifically deep learning methods have been developed that are able to capture far more complex binding motifs^5,6^. These methods have also been employed to predict variants likely to alter TFBSs and have thus become an essential component of many variant prioritization pipelines, such as the variant effect predictor^7^.

The performance by which TF binding models are able to distinguish their binding regions from random genomic regions has been well characterized^8,9^. To assess how well these predictors perform at identifying the impact of variants, known regulatory variants are often employed, which include variants from the Human Genome Mutation Database (HGMD), genome-wide association studies (GWAS), and quantitative trait loci (QTL) studies^10-12^. Yet, little has been done to explore the ability of these models to assess the impact of genetic variants on binding in a TF-specific manner. Allele-specific ChIP-seq binding (ASB) is a valuable resource to carry out such performance assessments. Here, ChIP-seq reads are mapped to either allele of heterozygous variants within an individual or cell line, allowing for the explicit identification of variants that do and do not alter TF occupancy. Several studies have utilized ASB data to explore TF-specific performance at assessing variant impact. For instance, Zeng et al. used ASB variants for six TFs to validate their GERV method at identifying TFBS-altering variants^13^. Shi et al. compiled a dataset of over 10,000 ASB variants across 45 ENCODE ChIP-Seq datasets and demonstrated that ASB variants lie within highly relevant PWM positions^14^. These studies are, however, often based on a small number of TFs or are focused on individual variant impact methods.

In this study, we aimed to carry out a systematic and unbiased analysis of the performance of TF-binding models at assessing variant impact. We compiled and collected a compendium of over 132,373 potential ASB variants across 101 TFs. We define ASB variants for 81 of these TFs and compare the performance of five different methods, based on PWMs, deep learning and k-mer-based machine learning, at predicting impactful variants through different measures of variant impact for each method. In some cases, these measures perform differently at variant impact prediction. We show that, overall, deep learning and k-mer-based machine learning methods significantly outperform that of the commonly-used PWMs. We also explore the performance of TFs individually, identifying TFs that are able to accurately predict variant impact as well as those that, although have distinct binding specificities, are unable to do so. We finally investigate mechanisms such as methylation, DNA shape, PTMs and co-binders that may explain poor performance in variant impact prediction. We show that ASB data serves as a valuable benchmarking resource on which the performance of TF binding models can be exhaustively surveyed with respect to variant impact.

## Results

### 1.1 A compendium of allele-specific binding events

To assess the performance of TF-binding impact predictions, we require a set of variants known to alter binding, whether the alteration is a gain or loss, and also a set of variants known to not alter binding. This is conveniently provided by ASB data. We collected ASB variants from five studies^14-18^, with each study providing heterozygous variants, the sample or cell line from which it was obtained, and reference and alternate allele read counts and the TF affected (**Figure 1a**). Since read counts were collected from different sources and processed with their respective analysis pipelines, we also ensured results from different studies were consistent with one another where possible. We correlated the effect sizes of overlapping ASB variants from different studies and found a high degree of concordance (mean Pearson r = 0.79, **Supplementary Figure S1**). In line with this reasoning, if an ASB variant was reported across multiple studies, we retained the entry with highest number of total mapped reads. We additionally discarded variants with fewer than 10 reads mapping to the reference or alternate allele, for a total of 132,373 variants and 215,631 TF-variant pairs (potential ASB events) reported across 101 TFs. The largest fraction of TFs was reported in a single study, with a total of 50 TFs and as few as three TFs were reported across all five studies (**Figure 1b**). Different studies also contained a disproportionate number of TFs and samples for which ASB data was available. The largest number of TFs was contained within the Santiago et al. dataset with a total of 80 TFs across from 14 samples^15^ (**Figure 1c-d**).

**Figure 1.**
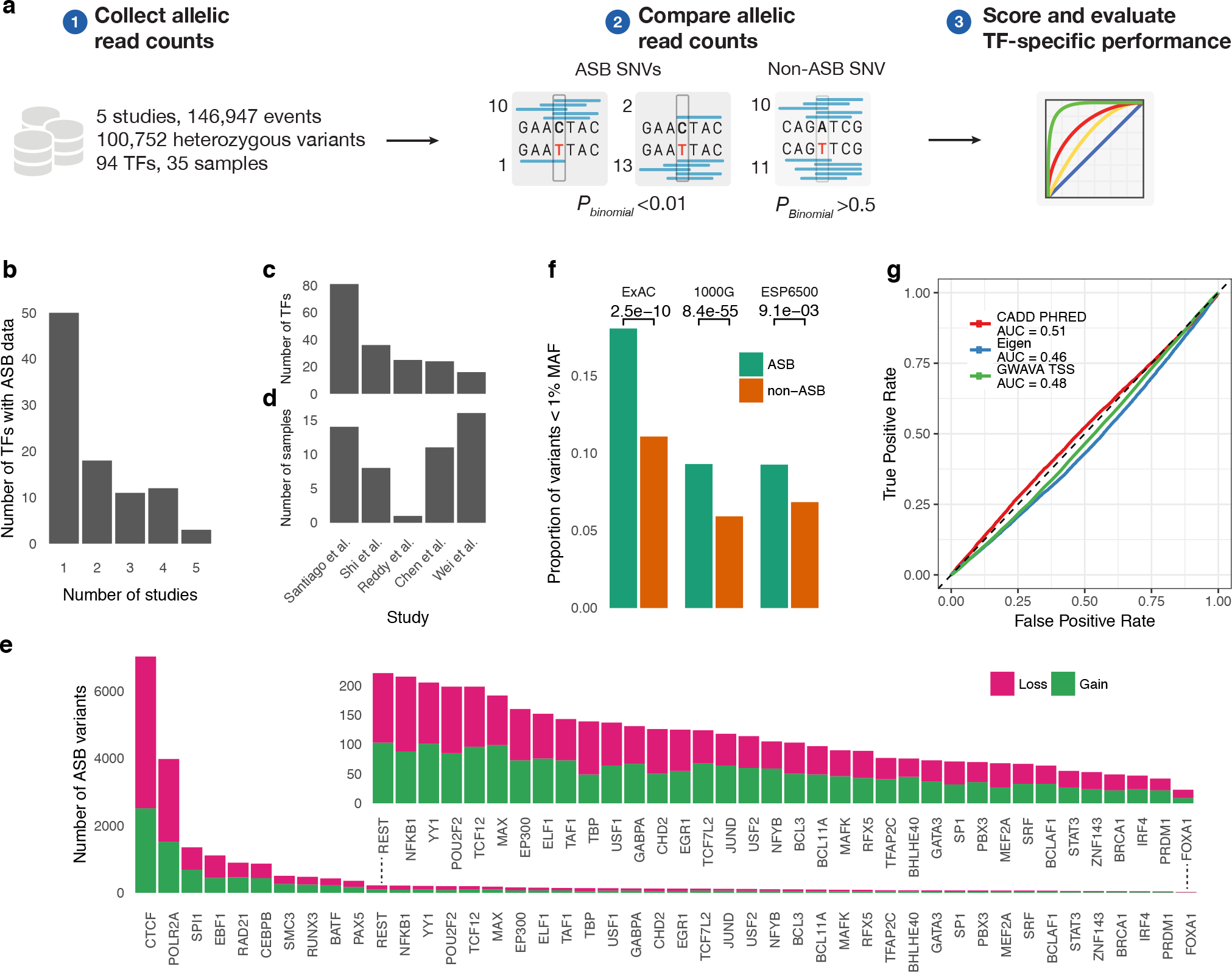
Properties of ASB and non-ASB variants. (a) The use of ASB data for assessing the performance of TFBS variant impact. (b) The total number of TFs covered by a different number of studies. Only three TFs have ASB data in all five studies. (c-d) The number of TFs and samples per ASB study. (e) The number of ASB variants per TF at a *P*_binomial_ < 0.01 and at least 10 reads mapped to either allele. Only TFs with at least 20 ASB variants are shown. Loss and gain ASB variants are shown in magenta and green, respectively. (f) ASB variants (green) are relatively rare compared to that of non-ASB variants (orange). Significance *p*-values represent a one-sided Fisher’s exact test. (g) Non-coding variant impact predictors are unable to distinguish ASB variants from non-ASB.

The binomial test was used to define how the significance of the imbalance between the reference and alternate read counts (**Methods**), which is commonly used in ASB studies^16,19^. ASB variants that exhibit significant differences between reference and alternate read counts were defined by a significance threshold *P*_binomial_ < 0.01, resulting in 32,252 ASB events across 81 TFs, of which 58.1% (18,744) were loss events where the alternate read count is lower and 9,397 (41.9%) were gain events, where the reference read count was lower. Although one would expect that there would be a significantly higher number of losses than gains, since disrupting a binding site is mechanistically easier, our results indicate that overall a random variant has just as much chance of creating a site as it does to destroy one. A total of 79,827 non-ASB events with balanced read counts were defined as those with *P*_binomial_ > 0.5 (**Methods**). Overall, 81 TFs had at least one defined ASB and non-ASB event, whereas 58 TFs had at least 10 events and 51 TFs had at least 20 events. To our knowledge, this is the largest available ASB dataset curated.

ASB variants are implicated in altering TF-binding and should be less likely to exist with high frequency. We confirmed this by analysing the proportion of ASB variants which are rare at a MAF <1% using data from the ExAC consortium^20^, 1000 genomes project^21^, and the ESP6500 project^22^ (**Methods**). Strikingly, loss ASB variants consistently demonstrated a higher fraction of rare variants, compared to non-ASB variants (*p*<1.57×10^−9^), which was in contrast to gain ASB variants that did not show any significant enrichment (*p*<0.98, **Figure 1f**). This suggests that loss of binding would be more likely consequential, which is similar to what had been previously observed for whole genome *de novo* variants in autism^23^. We additionally assessed whether commonly used non-coding variant impact predictors, such as GWAVA^24^, Eigen^25^ and CADD^26^, could accurately distinguish ASB variants from non-ASB using the area under the receiver operating characteristic curve (AUROC) measure (**Methods**). However, near-random performance was observed for all three methods (CADD AUROC = 0.51, Eigen AUROC = 0.46, GWAVA AUROC = 0.48, **Figure 1g**). This suggests that current approaches, which do not incorporate TF specificity are unable to identify variants altering TF-binding.

We utilized the collected ASB data to assess and compare the performance of several computational predictors of TF-binding variant impact (**Figure 1a**). The approaches included in the analysis were those based on PWMs^1, 5^, *k*-mer-based approaches GERV^13^ and gkmSVM^12^, and deep learning-based approaches DeepBind^5^ and DeepSEA^6^.

### 1.2 Scoring metrics for evaluation of transcription factor binding variant impact

The different methods available offer a variety of scoring metrics that describe the quantitative impact of a variant on TF-binding. These metrics are typically signed, where strong negative and positive values indicate loss and gain, respectively. DeepSEA produces a single probability of binding for both the wildtype and mutant sequences and uses two metrics to quantify the impact of a variant: the difference (*diff*) and log fold change (*log FC*) between the probabilities (**Methods**). gkmSVM provides a single score *deltaSVM* reflecting the change in the sum of k-mer weights for wildtype and variant sequences and GERV provides a single unsigned score (*GERV score*) that reflects the change in predicted ChIP-seq read counts (**Methods**).

In contrast, PWMs and DeepBind only provide a score reflecting the likelihood of binding and not the impact of a variant. For these approaches, we devise a number of metrics to assess the impact of a variant. Because the TF specificity models receive as input a fixed length sequence, which may not correspond to the length of a sequence underlying an ASB event, we score multiple overlapping fixed-length windows of sequences along the region of interest with the reference and alternate allele. The defined metrics serve as a good starting point for assessing how different approaches perform at scoring the variant impact on TF-binding and are described in more detail.

#### 1.2.1 Raw score metrics of variant impact

The difference in raw model scores is typically used to assess the impact of a variant^27-29^ (**Supplementary Figure S2**). We similarly define *delta raw* as the maximum difference between the raw wildtype and mutant scores across the *k*-mers. Because TF-binding can be made robust through homotypic clusters of redundant binding sites, they can often mitigate effects of impactful variants^30^. In line with this reasoning, we devised *delta track* as the difference between the maximum of all wildtype window scores and the maximum of all mutant window scores. Both metrics are signed, such that losses are indicated by negative scores and gains by positive (**Methods**).

#### 1.2.2 Probability-transformed metrics of variant impact

We sought to aid interpretability and strengthen baselines for variant effect prediction. To do this, we convert raw scores (which are not on any particular scale and not comparable across TFs) to likelihoods of binding (which are normalized to [-1, 1] and are comparable across TFs). We define positive and negative sequences as those used to train the DeepBind or PWM model and random genomic regions, respectively (**Methods**). We found that, particularly for ChIP-seq/SELEX data used to train DeepBind models, distributions of raw scores from the background followed a normal distribution and in some cases, distributions of foreground scores were bimodal with one component of scores exhibiting similar properties to that of the negative distribution (**Supplementary Figure S3**). This is likely due to lenient threshold used to call the ChIP-seq peaks, which was done to maximise the number of sequences available to train models. Since the foreground distribution is bimodal, we use a Gaussian mixture model (GMM) to learn the two components comprising the foreground distribution. One component is fixed to the parameters of the negative distribution and the “true positive” component is learned. A linear model is then trained and used to compute the probability of binding (*P_bind_*) from raw PWM or DeepBind scores (**Methods**).

Using the *P_bind_* score, we define the *delta P_bind_* score as the maximum difference between the mutant and wildtype *P_bind_* probabilities across all sequence windows. This value ranges from −1 to 1, where low negative values indicate a loss of binding and high positive values indicate a gain of binding.

We additionally define a probabilistic score *P_loss_* and *P_gain_* that range from 0-1 reflecting the likelihood of a binding site being lost or gained, respectively. For *P_loss_,* this is computed by taking the joint probability of binding for the wildtype sequence and the probability of the mutant not binding and vice versa for *P_gain_* (**Supplementary Figure S2, Methods**). *P_loss_* and *P_gain_* are combined into a single score by first signing *P_loss_* negatively and computing the probability with the higher absolute value as *P_comb_*. Since *P_loss_* and *P_gain_* can have low to moderate magnitudes we also compute *P_sum_* as the sum of the signed probabilities, resulting in a near-zero score for such cases (**Methods**).

### 1.3 The use of allele-specific binding data for benchmarking variant impact prediction

Given the numerous available predictors and scoring metrics available for prioritising the impact of variants on TFBSs, we investigated how well each method and scoring metric performed at distinguishing TFBS-altering variants using the ASB data as a gold standard.

We collected and trained models for TFs with ASB data from the described methods. Pre-trained DeepBind models for 91 TFs were utilised, which we had previously trained on ENCODE ChIP-seq data^5^. For DeepSEA, pre-trained models for 91 TFs were used that were trained on similar ChIP-seq datasets to those used for DeepBind, matched by the cell line from which the training data was obtained. The same data used to train DeepBind models was used to train 91 corresponding gkmSVM models and pre-trained GERV models for 60 TFs were collected, based on ChIP-seq data (**Methods**). We further utilised PWMs for 56 TFs from the JASPAR database along with 87 sets of PWMs based on over-represented motifs discovered by MEME-ChIP^31^ from the data used to train DeepBind models. For each TF, sequences matching a set of the top five over-enriched motifs were used to construct at most five PWMs. Using the set of PWMs, predictions were generated for the (1) “signif” most significant PWM and (2) “best” the PWM that resulted in highest magnitude variant-impact score (**Methods**).

Each method, model, and scoring metric was used to score both ASB and non-ASB data. The resulting scores were used to assess the performance of the predictor at discriminating variants implicated in loss or gain ASB from that of non-ASB variants using the receiver operator curve (ROC) and precision-recall (PR) curve. The AUROC and area under PR curve (AUPRC) are used to provide a quantitative measure of performance, where both metrics provide a different view on performance.

The performance was measured using the definition of an ASB and non-ASB variants as those with a *P*_binomial_ < 0.01 and *P*_binomial_ > 0.5, respectively and exhibited at least 10 reads mapped to the reference or alternate allele. We further only considered 51 TFs with at least 20 ASB and non-ASB variants to improve robustness. Of these TFs, 82% (42/51) had a DeepBind, DeepSEA or gkmSVM model, 73% (37/51) had a GERV model, 55% (28/51) had a JASPAR PWM model and 76% (39/51) had a MEME-ChIP-based PWM model.

For DeepBind, we found that ChIP-seq models generally outperform that of SELEX at predicting ASB variants with respect to the AUROC (*p*<0.028, **Supplementary Figure S4**). Certain TFs such as USF1, IRF4 and CTCF showed notably improved performance with ChIP-seq models (**Supplementary Figure S5**). Furthermore, DeepSEA models were only trained on ChIP-seq models. As a result, we discarded SELEX models from the analysis.

#### 1.3.1 A comparison of variant-impact scoring metrics

We first explore performance of PWM-based scoring metrics. We compared the performance of five scoring metrics used for PWMs in JASPAR and MEME-based PWMs. The performance of scoring metrics in most cases was equivalent to one another in each of the PWM sets, with the exception of *delta raw* which consistently demonstrated poor performance (*p*<8.5×10^−4^, **Supplementary Figure S6**). For instance, in JASPAR PWMs, the average AUROCs for *delta raw* and *delta track* was 0.58 and 0.53 (Δ*AUROC* = 0.051), respectively and 68% (19/28) of TFs showed a 10% increase in *delta track* performance for either loss or gain ASB events (**Figure 2a**). By inspecting failure cases, we observed that *delta raw* was often incorrect due to the inflation of scores caused by maximising differences overall sequence windows. High *delta raw* scores do not necessarily indicate a loss or gain of due to the positional independence of PWMs. For instance, a low-scoring wildtype sequence harbouring a variant in a position of importance for the PWM will result in a high *delta raw* score. This effect coupled with taking the maximum over sliding sequence windows results in an inflation of scores, which affects the identification of true negatives (non-ASBs) and true positives (ASBs). These effects are only partially mitigated by metrics such as the *delta track* and probabilistic metrics (**Supplementary Figure S6**).

**Figure 2.**
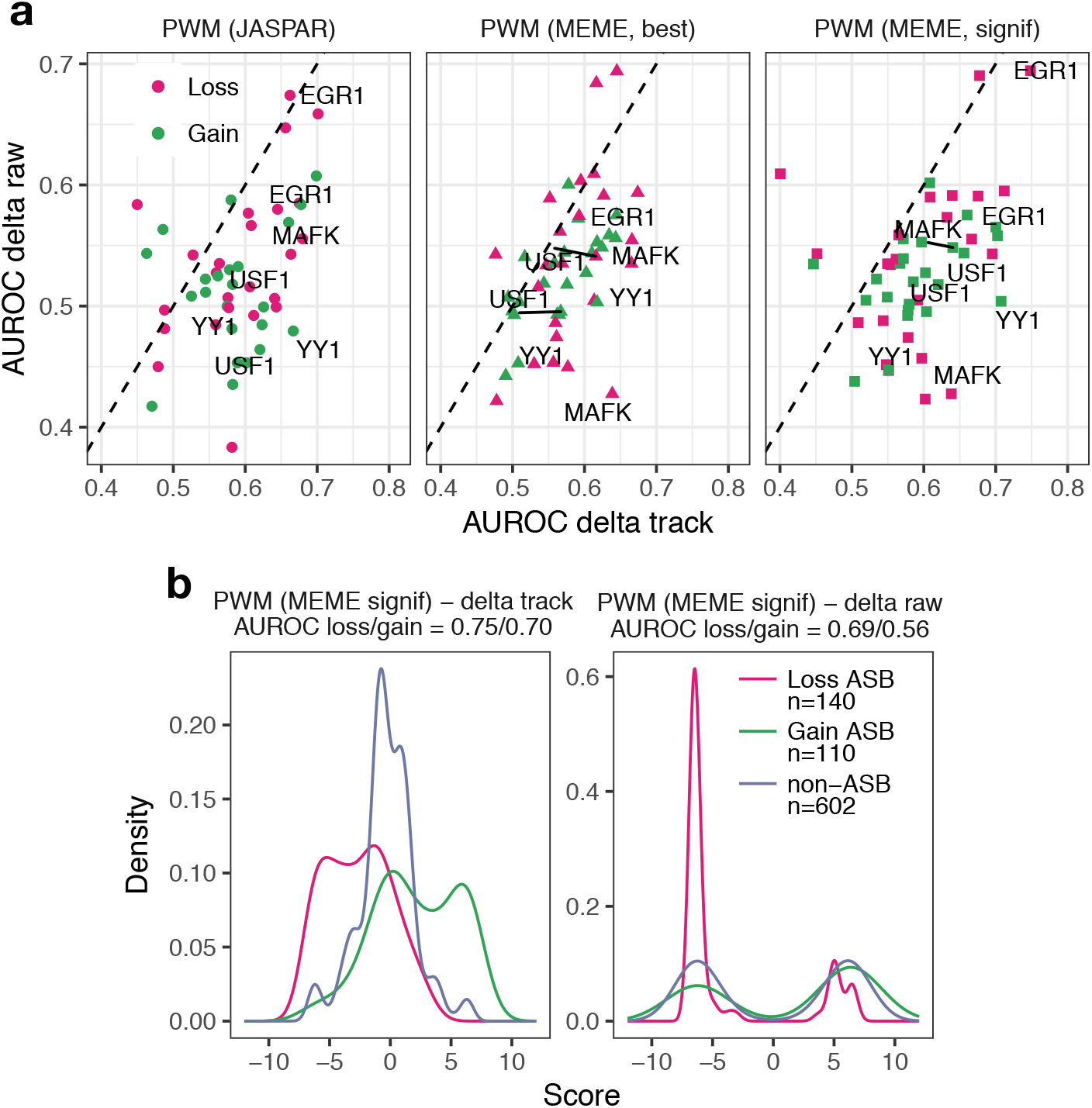
Comparison of *delta raw* and *delta track* metrics for the PWM. (a) AUROCs for *delta track* compared to that of *delta raw* for the three PWM sets. (b) Density plots showing the inflation of scores in the *delta raw* metrics for the EGR1 TF.

The early B-Cell Factor 1 (EBF1) is one of the TFs with lower performance for *delta raw* compared to the other metrics. For the MEME signif PWM, the *delta track* showed an AUROC of 0.75 and 0.70 (Δ*AUROC* = 0.05) for loss and gain, respectively whereas *delta raw* showed AUROCs of 0.66 and 0.61 (Δ*AUROC* = 0.05), respectively. **Figure 2b** shows the distributions of scores for loss and gain ASB and non-ASB variants, highlighting the inflation of scores for non-ASB variants.

Detailed examples showing the calculation of *delta raw* and *delta track* for an EBF1 ASB and non-ASB variants are shown in **Figure 3a-c**. Here, the scores for the wildtype and mutant track are shown, along with the difference for each sequence window and the final computed scores. The first example highlights a non-ASB variant, where a near-zero predicted score is desired, yet despite no predicted binding occurring on either the wildtype or mutant tracks, the *delta raw* metric still results in an inflated score through differences computed in non-binding regions (**Figure 3a**). The second example highlights a loss ASB variant and the loss event is correctly identified by both metrics. However, the maximum difference for *delta raw* here is obtained not from the sequence window exhibiting the loss (window 19), but rather at another window 14) (**Figure 3b**). The third and final example highlights a gain ASB event that shows how the drawbacks of the *delta raw* metric lead to an incorrect prediction of the variant as loss, whereas *delta track* correctly predicts the directionality (**Figure 3c**). The *delta track* and similar metrics offer numerous advantages over identifying the largest possible difference. All metrics will, however, be bottlenecked by the high degree of false positives produced by PWMs.

**Figure 3.**
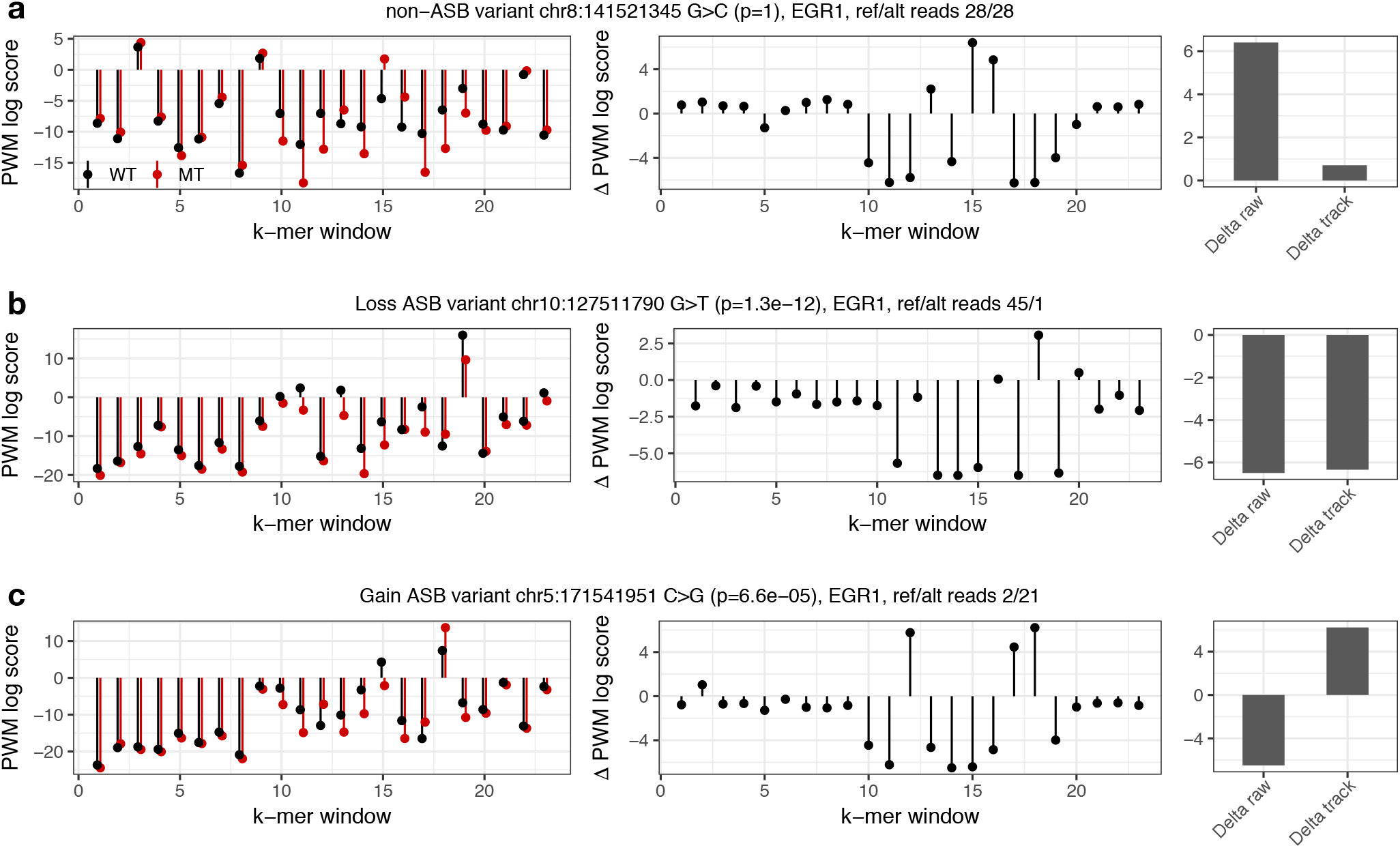
EGR1 examples of scores for individual sequence windows highlighting the differences between *delta track* and *delta raw*. The left plot shows the wildtype (black) and mutant (red) scores for each window, the middle plot shows the score difference, and the right plot shows the final *delta raw* and *delta track* scores. This is shown for (a) a non-ASB that are misclassified by *delta raw,* and correctly classified by *delta track* (b) loss ASB correctly identified by both metrics, and (c) gain ASB misclassified by *delta raw* and correctly identified by *delta track*.

The performance of scoring metrics used in both DeepBind and DeepSEA were also assessed. We compared the two DeepSEA metrics and found that, overall, neither metric significantly outperformed the other (*p*<0.42). The *log FC* metric did, however, show an average AUROC increase of 0.03 (mean AUROC 0.62 vs. 0.59) and 0.019 (mean AUROC 0.64 vs. 0.62) for loss and gain, respectively (**Supplementary Figure S8**a). For DeepBind, no significant difference was observed in performance between the five used metrics. Unlike the PWM, the *delta raw* did not overall show difference to that of *delta track* (Δ*AUROC* < 0.009).

The choice of the scoring metric used in variant impact can often be critical to both interpretability and performance. For PWMs, the *delta raw* metric in PWMs has been long used in studies to quantify effect of a variant of a TFBSs^27-29,32^. The results demonstrated here indicate that the choice of score metric when using PWMs, particularly *delta raw,* can drastically impact the reliability of predictions made on regulatory variants. Alternative metrics such as *delta track* and the probabilistic metrics *P_comb_* and *P_sum_* offer good approaches to mitigating effects by *delta raw* but still are bottlenecked by the inherent limitations of PWMs. For deep learning approaches, little overall difference was observed between metrics and the choice of metric in this case remains purely for interpretation purposes.

#### 1.3.2 Performance of binding models vary depending on the definition of ASB variants

We then asked whether performance varied if thresholds used to define ASB and non-ASB variants were changed. We measured the AUROC for a combination of thresholds for both the *P*_binomial_ (*p* < 0.1, 0.01, 10^−3^,10^−4^and 10^−5^) and the minimum number of reference or alternate reads (≥10, ≥20 and ≥30 reads). Performance was measured for seven TFs (BATF, CEBPB, CTCF, EBF1, RUNX3, SMC3, TBP) which had ≥20 ASB variants at 10^−5^ and ≥30 reads.

Utilising performance measures for seven TFs with sufficient data at ≥ 10 reads and *P*_binomial_ < 10^−5^ we found that, on average, more stringent definitions of *P*_binomial_ thresholds exhibited higher AUROCs, which was consistent across both loss and gain ASBs (**Supplementary Figure S7**a). For instance, with gain ASBs at ≥10 reads, DeepBind had an average AUROC of 0.70 and 0.59 at *P*_binomial_ < 10^−^5 and *P*_binomial_ < 0.10 thresholds, respectively. Conversely, increasing the minimum number of reads did not show any substantial shift in performance (**Supplementary Figure S7**a). These results suggest that models are better able to distinguish variants with a higher imbalance in the number of reads and that higher read imbalance is more likely driven by changes in sequence specificity.

To determine if more stringent thresholds used to define non-ASB variants affected the AUROC, we fixed the ASB *P*_binomial_ to 0.01 with ≥ 10 reads and assessed performance at different *P*_binomial_ thresholds of 0.5, 0.7, and 0.9 for 21 TFs which had at least ≥20 non-ASB variants at *p* > 0.9. However, higher thresholds of *P*_binomial_ did not show any significant variation in performance (**Supplementary Figure S7**b).

More stringent thresholds of *P*_binomial_ result in modest increases in performance. This, however, comes at a cost of losing a majority of TFs for which ASB data is available (11 TFs at *P*_binomial_ < 10^−5^). Thus, to assess performance with a sufficient number of TFs, we retain the thresholds of *P*_binomial_ < 0.01. Furthermore, since no significant increase was observed at higher reads we retain the ≥10 reads for further analyses. It is noteworthy to mention that at the threshold of *P*_binomial_ < 0.01 there still exists a reasonable imbalance in read counts (for instance, 100 vs. 37 read counts would yield a *P*_binomial_ = 0.012). While parts of this may be noise, we expect that many include ASB events that impact TF occupancy through the alternative mechanisms described and are not detected by current *in silico* models. Lastly, it is not expected that the performance of gain and loss differ significantly and any changes observed are likely due to the small number of TFs being assessed.

#### 1.3.3 Machine learning-based methods outperform PWMs at predicting the impact of variants on transcription factor binding

PWMs have been the *de facto* approach to modelling TF specificity and assessing the impact of regulatory variants on TF-binding. Machine learning, and in particular deep learning approaches are able to capture more complex relationships and reduce false positive predictions. We, therefore, next asked how the performance of machine learning *k*-mer-based as well as deep learning approaches compared to that of PWMs at predicting the variant impact. For methods with more than one scoring metric, we selected the top performing metric, which included *delta track* for PWMs, *log FC* for DeepSEA and *delta raw* for DeepBind, the *deltaSVM* score from gkmSVM and the *GERV score* from GERV. We compared performance based on the AUROC and AUPRC (**Figure 4**).

**Figure 4.**
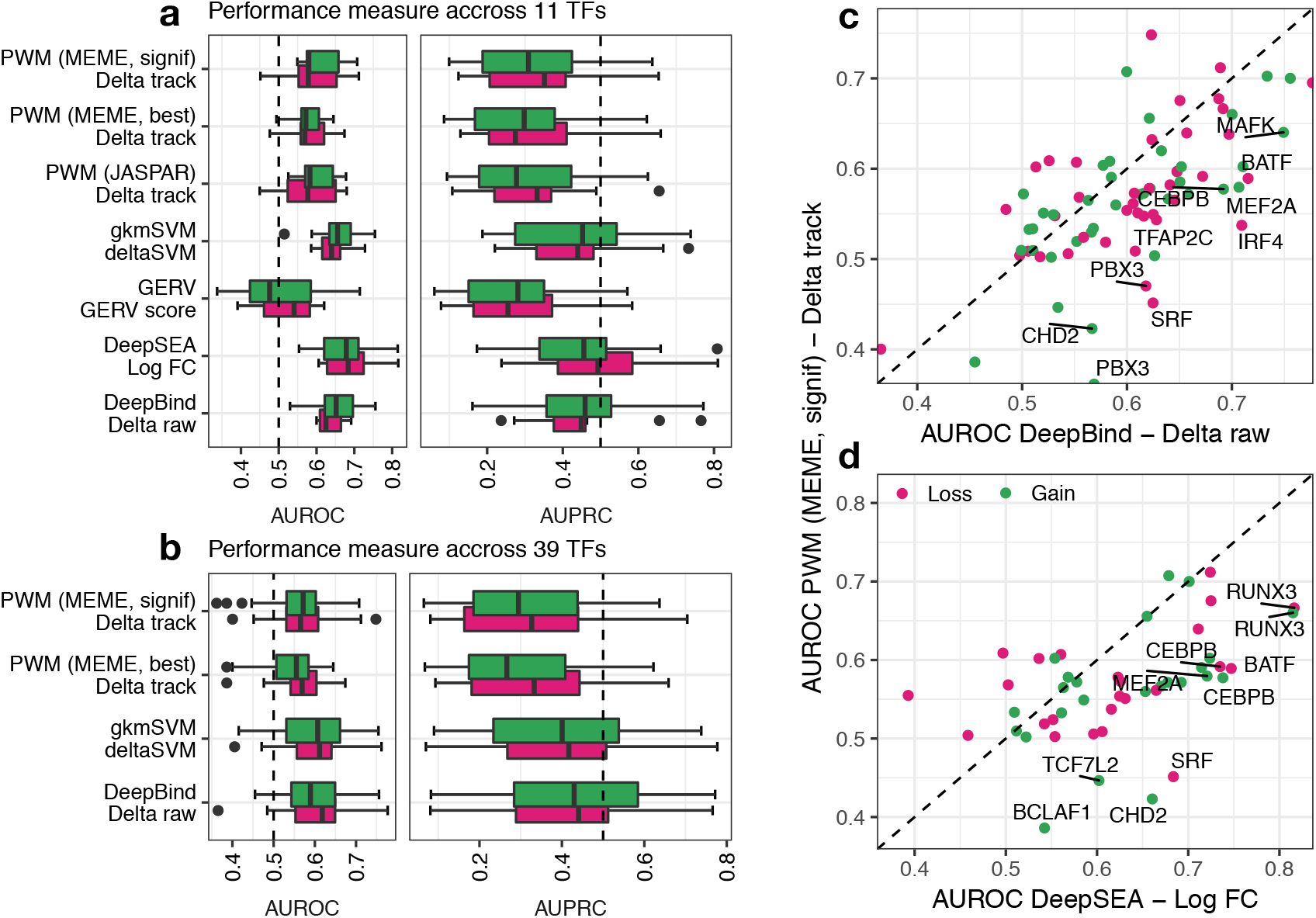
Comparing performance of machine learning approaches to PWMs. (a-b) Comparison of AUROCs (left) and AUPRCs (right). Performance is shown for (a) 11 TF models shared amongst five methods and for (b) 39 TF models shared amongst four methods. (c-d) Scatter plots showing the AUROCs for individual TFs for deep learning models (c) DeepBind *delta raw* and (d) DeepSEA *log FC* against PWM *delta track*.

Because there exists a different number of trained models with ASB data for each method, we compared performances for 11 TFs with models across all five methods. GERV showed near random performance across both loss and gain, performing poorer than PWMs. The other machine-learning approaches including gkmSVM, DeepBind and DeepSEA significantly outperformed PWMs with respect to AUROCs (**Figure 4a**, *p*=0.034 DeepBind, *p*=5.91 × 10^−04^ DeepSEA, *p*=0.038 gkmSVM) and AUPRCs (p=4.57 × 10^−3^ DeepBind,*p*=1.24 × 10^−3^ DeepSEA,*p*=0.024 gkmSVM). We further limited the predictors being compared in order to retain a larger number of common models between the methods. We compared MEME-based PWMs, with gkmSVM and DeepBind for a total of 39 common TFs, where both DeepBind and gkmSVM similarly outperformed the PWM-based models with respect to AUROCs (**Figure 4b**,*p*=4.07 × 10^−3^ DeepBind,*p*=4.85 × 10^−3^ gkmSVM) and AUPRCs (*p*=2.32×10^−3^ DeepBind, *p*=7.31×10^−3^ gkmSVM).

Comparing the AUROC of deep-learning-based methods to that of PWM *delta track,* we identify the TFs SRF, CHD2, IRF4, BATF and CEBPB amongst those where deep learning models perform better at predicting variant impact (**Figure 4c-d**). A more in-depth examination of DeepBind and PWM scores reveals that even the best performing PWM metric often results in high numbers of false positives and false negatives. These results further illuminate the importance of machine learning models in variant impact.

#### 1.3.4 Alternative binding mechanisms explain differences in variant impact prediction performance

Having established that machine learning approaches outperform PWMs, we sought to focus on DeepBind and DeepSEA models and investigate the performance of individual TFs.

Amongst TFs that performed well are RUNX3, BATF, MAFK, and CEBPB, which had an AUROCs of > 0.7 in either DeepBind or DeepSEA models. Conversely, TFs like SP1, BRCA1, TBP, and TAF1 consistently showed near-random performance (**Figure 5a-b**). We asked whether poor ASB performance is dictated by the model’s performance at identifying binding sites. A model that is unable to correctly identify binding sites should not perform well at identifying the impact of ASB variants. Indeed, we found that models with an AUC < 0.80 in DeepBind also demonstrated poor ASB performance (**Figure 5c-d**). However, high-performing models showed a high degree of variation with respect to ASB performance. For instance, SP1 and BRCA1 both have model AUROCs of 0.99. Despite having explicit sequence specificities, such models are unable to detect *in vivo* occupancy differences introduced by variants. This suggests alternate mechanisms beyond simple binding site specificities affecting binding. Since a number of mechanisms have been shown to contribute to TF-binding specificities such as methylation, DNA shape, TF cofactors and regulatory PTMs on the TF, we explored whether such mechanisms are able to explain the poor performance observed.

**Figure 5.**
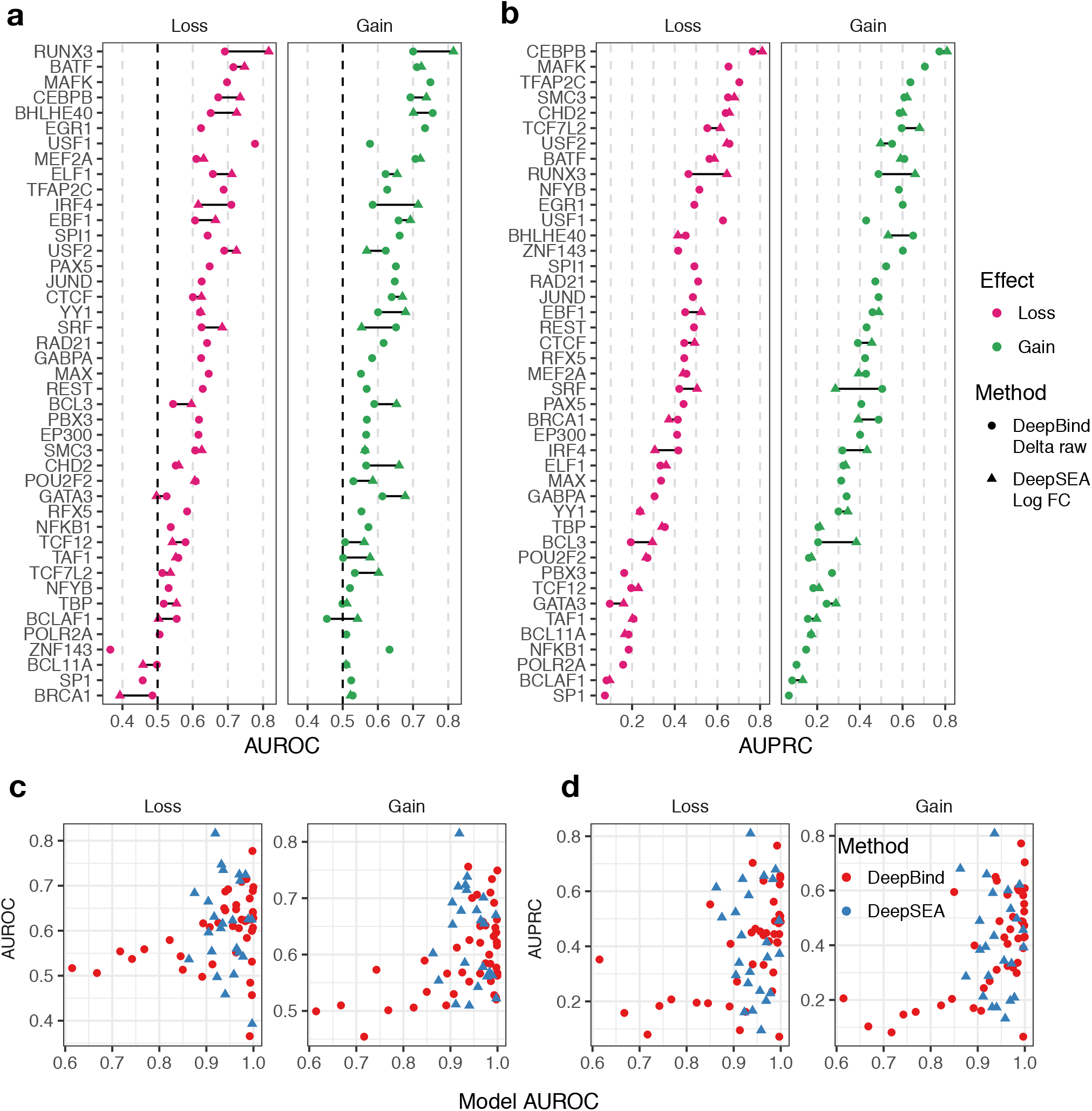
Exploring performance of individual TFs for deep learning methods. (a) AUROCs and (b) AUPRCs for loss (magenta) and gain (green) ASBs. TFs are ordered by the maximum performance metric across methods and effects (c-d) AUROCs of binding performance is compared against performance of models to identify impact of variants, as defined by (c) AUROCs and (d) AUPRCs for DeepBind *delta raw* (red) and DeepSEA *log FC* (blue) models.

TFs that are involved in binding complexes can obtain their specificity by the binding of cofactors^33^. We collected known physical TF-TF interactions from the transcription cofactors (TcoFs) database^34^ for 35 TFs with performance measures and asked whether the degree of interactors predicted ASB performance. We found that TFs such as the TATA-binding protein (TBP) and the specificity protein 1 (SP1) which showed upwards of 50 interactions with other TFs also correlated with poor performance (**Figure 6a**).

**Figure 6.**
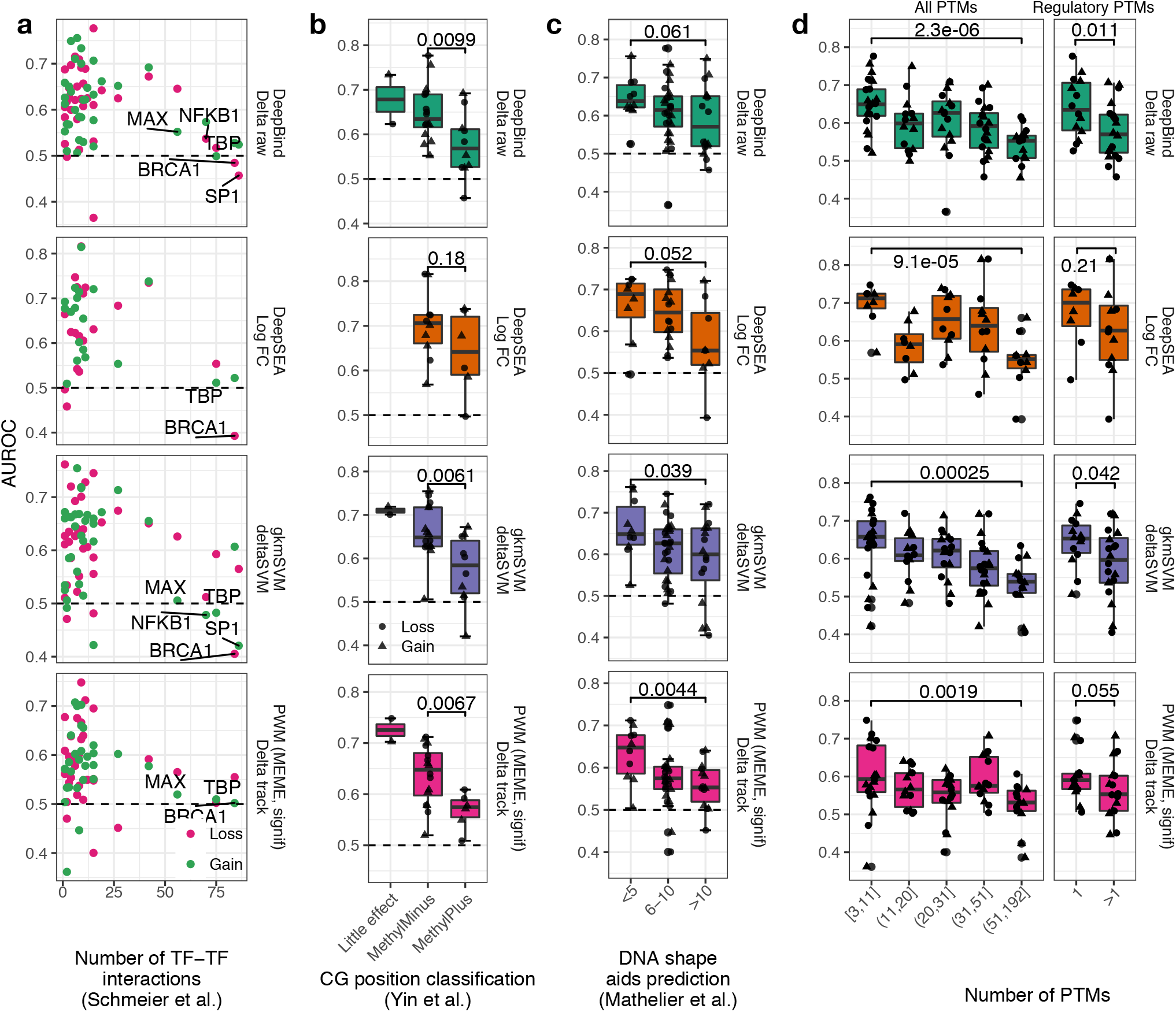
Alternative mechanisms that contribute to poor variant-impact prediction. Performance, as measured by AUROCs across DeepBind, DeepSEA, gkmSVM and PWMs for (a) Number of TF-TF interactions, (b) MethylPlus versus MethylMinus TFs (c) TFs where binding is influenced by DNA shape and (d) PTMs. Significance *p*-values are based on a one-sided Wilcoxon test.

Cytosine methylation is another major factor shown to dictate binding for many TFs^35^. A recent study by Yin et al. used a methylation-sensitive derivative of SELEX to identify TFs influenced by methylation for over 500 TFs^35^. Here, TFs were characterised into three groups MethylPlus, where the TF preferred to bind methylated sequences, MethylMinus, where little to no TF binding was found for methylated binding sites and LittleEffect, where methylation had little to no effect on binding. We collected classes for 14 TFs with performance measures across the different methods and compared performance for each class. Interestingly, we found that TFs classified as MethylPlus consistently showed significantly lower performance compared to that of MethylMinus in DeepBind (**Figure 6b**, *p*=9.9×10^−3^), gkmSVM (*p*=6.1 × 10^−3^) and PWMs (*p*=2.8 × 10^−4^) (**Figure 6b**). MethylPlus TFs included included SP1, RFX5, POU2F2, and GATA3, which demonstrated low average AUROCs across methods RFX3 (0.53), SP1 (0.49), POU2F2 (0.55), GATA3 (0.59). Indeed, TFs such as RFX3 and SP1 methylation has been shown to positively regulate binding^36,37^. This suggests that TFs relying on methylation for binding are likely to perform worse when only sequence information is used for model training.

TFs are known to be able to detect three-dimensional shape of DNA^38^. We utilised data from Mathelier et al., where models were trained that incorporated DNA shape features to show performance of binding can be improved^39^. We identify TFs that rely on DNA shape for binding by computing the percent increase in AUROC (Δ%) for models. The percent increase is binned values into three bins, < 5,6 - 10, and > 10, which represent minimal improvement, medium improvement and strong improvement. We found that TFs that showed strong improvement had significantly lower performance when compared to those that showed minimal improvement for PWMs (*p*=4.4×10^−3^) and gkmSVM (*p*=0.039). Significance was borderline significant for DeepBind (*p*=0.061) and DeepSEA (*p*=0.035). This can be explained by the fact that deep learning approaches should at least in part be able to extract DNA shape features from the sequences they are trained on (**Figure 6c**). TFs in which DNA shape aided binding prediction (>10%) included SRF (mean AUROC = 0.62, Δ% = 23), BRCA1 (mean AUROC = 0.48, Δ% = 15), NFYB (mean AUROC = 0.50, Δ% = 12.6), MEF2A (mean AUROC = 0.62, Δ% = 12), MAFK (mean AUROC = 0.68, Δ% = 11.7), TBP (mean AUROC = 0.53, Δ% = 11), PAX5 (mean AUROC = 0.63, Δ% = 10.1), and SP1 (mean AUROC = 0.50, Δ% = 10.1). In contrast, TFs where DNA shape played a minimal role in binding prediction included ELF1 (mean AUROC = 0.65, Δ% = 1.8), TFAP2C (mean AUROC = 0.62, Δ% = 2.6), BHLHE40 (mean AUROC = 0.69, Δ% = 3.5), and USF2 (mean AUROC = 0.65, Δ% = 3.8). This suggests DNA shape as a valuable feature when assessing variant impact on TF-binding.

Finally, we explore the impact of PTMs on TFs binding. PTMs are key regulators of transcriptional activity and are known to govern binding specificity^40^. We asked if TFs that are more likely regulated by PTMs also performed poorly. We collected 1,645 PTM sites for seven modifications in 43 TFs from PhosphoSitePlus^41^. We binned the TFs by the number of PTM sites they harboured by percentiles. In many cases, we found that heavily modified TFs such as BCLAF1 (mean AUROC = 0.49, n = 192), POLR2A (mean AUROC = 0.52, n = 171), EP300 (mean AUROC = 0.59, n = 139) and SMC3 (mean AUROC = 0.57, n = 90) showed significantly lower performance levels, compared to TFs that harboured fewer than 10 PTM sites in DeepBind (*p*=6.6×10^−5^), DeepSEA (*p*=5.6×10^−5^), gkmSVM (*p*=2.1×10^−4^) and PWMs (*p*=1.6×10^−3^, **Figure 6d**). We similarly utilised 42 PTM sites across 17 TFs known to be regulatory and compared performance of TFs with a single regulatory PTM to those with more than one. We observed a similar trend, where TFs such as BRCA1 (mean AUROC = 0.48, n = 6), SP1 (mean AUROC = 0.50, n = 5), and NFKB1 (mean AUROC = 0.51, n = 4) with a high number of known regulatory PTMs displayed significantly lower performance across DeepBind (*p*=0.011) and gkmSVM (*p*=0.042) **Figure 6d**).

For certain TFs with distinct sequence specificities, elucidating the impact of variants can be more challenging due to the sequence specificity depending on a multitude of factors that involve mechanisms beyond proximal sequence information alone. These results demonstrate the importance of such factors, *in silico,* when assessing variant impact in TFBSs.

## Discussion

Understanding the impact of non-coding variation is an ongoing challenge in genetics. One of the primary modes for this is through impacting TFBSs. Yet, despite the wealth of TF specificity available through high throughput technologies, accurate *in silico* prediction of TFBS-altering variants remains a non-trivial task.

This study describes efforts to compare TF-based variant impact predictors using ASB variants as a gold standard. Since both alleles exist in the same cellular environment, ASB variants serve as a valuable source to assess the performance of TF-binding models at assessing the impact of variants. We have shown that the ability for machine learning models, in particular, deep learning methods, to significantly reduce the number of false positives allows for more accurate variant impact predictions. Deep learning approaches are able to utilise the full extent of ChIP-seq and SELEX data to learn far more complex positional dependencies in binding sites. Deep learning approaches are also not confined to the exact motif location and therefore can model sequence context of the binding site, which has been shown to contribute to binding^42,43^. We finally show that TFs with poor performance at assessing variant impact often rely on additional mechanisms such as binding partners, methylation, DNA shape and PTMs (**Figure 6**).

Assessing TF-DNA binding and how it is influenced by genetic variation *in silico* is a much more complex process than once thought. Current methods available for interpreting effects of TFBS variants rely primarily on binding specificity. Although this provides a useful framework for prioritizing non-coding variants, as demonstrated by results, even the most sophisticated methods are often unable to capture the full extent of altered binding in the genome. This can be attributed to several reasons. First, there are several other mechanisms that have been known to significantly contribute to binding specificities such as epigenetic modifications, cooperative binding, geometric shapes of DNA, PTM modifications and more. Epigenetics, in particular, methylation, can play a major role in enhancing or inhibiting TF-binding^35,44^. The recent study by Yin et al. carried out methylation-sensitive SELEX in 542 TFs and identified many methylation-dependent TFs^35^. Epigenetics can also greatly affect regions TFs can occupy. Nucleosome occupancy, for instance, results in closed chromatin which is inaccessible to TFs. Indeed, DNAase hypersensitivity sequencing (DHS-seq) has revealed regions of open chromatin in many cell lines, which have been shown to improve binding prediction^45^. PTMs is another major regulator of TF activity through altering its structural conformation, stability or sub-cellular localization thereby affecting binding^40^. For instance, phosphorylation of p53 on S378 allows it be recognized by 14-3-3 proteins, which associate with p53 and significantly enhances DNA-binding^46^. Second, binding preferences of a TFs have been shown to be heterogeneous across different cell lines. For instance, Arvey et al. comprehensively analysed ChIP-seq data for 67 TFs across multiple different and found that many cell-type-specific sequence models were able to capture binding variability, which was primarily due to differences in heteromeric complex formations^47^. Since the samples and cell lines from which ASB variants were obtained do not always match that of the experiments used to generate TF-specificity models, this is a potential confounding factor of poor performing models.

Another factor greatly limiting the prediction of TFBS-altering variants is the availability of TF motifs. It is estimated that the human genome contains approximately 1,400 TFs containing DNA-binding domains^48^. Although the current catalogue of TF-binding specificity has significantly expanded in the past decade with the aid of high throughput approaches such as ChIP-seq, SELEX and PBMs, almost half of identified TFs are yet to have their specificity determined^49^. This is perhaps due to technical limitations, such as transient binding or expression of the TF. The lack of such data further hampers ability for us to systematically understand variant impact in TFBSs.

Significant advances in interpreting non-coding variation have been greatly aided by the emergence of deep learning methods to the field of genetics over the past few years. However, accurate assessment of variant impact on TFBSs will require models to systematically integrate additional epigenetic, proteomic and genetic data in order to account for mechanisms beyond sequence specificity in a cell-type-specific manner.

## Methods

### 1.4 Collection of allele-specific binding data

ASB data were collected from five studies^14-18^. In each study, ChIP-seq reads are mapped to both alleles of heterozygous variants in individuals or cell lines. A count for the number of reads mapping to the maternal and paternal allele of each locus is provided by the studies. Allelic read imbalance is computed across all studies using a binomial test:

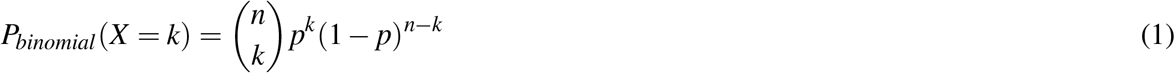

where *n* is the total number of reads mapped at a given loci, *p* is the probability of success, which is fixed to 0.5. This assesses deviation from the expected 50/50 read count. Finally, SNP positions for hg18-mapped variants are converted to hg19 using `liftOver`^50^ and loci that did not map were discarded. **Supplemntary Data 1** contains the complete list of curated ASB data utilised in this study.

### 1.5 Transcription factor binding model training and scoring

DeepBind models for a total of 91 TFs based ChIP-seq datasets were obtained from Alipanahi et al.^5^. Performance of each model was evaluated by applying models to left out test sequences (sequences not used to train the model) and random genomic regions. In the cases where there were multiple DeepBind models per TF, the model with the highest performance was selected. Scoring was carried out using the deepbind executable v0.11 with default parameters. Scores for DeepSEA were obtained through the online web server http://deepsea.princeton.edu/. Models for a total of 91 TFs were used that matched DeepBind models by cell line.

A total of 54 PFMs for 54 TFs with a DeepBind model were collected from JASPAR^1^. If a TF has more than one model, the model with the latest accession version is used. Motif enrichment data carried out using MEME-ChIP^31^ on ChIP-seq data used to train DeepBind models was used to construct a second set of PWMs. Each contained a set of enriched motifs along with matching ChIP-seq sequences and an *e*-value reflecting the enrichment significance. Motifs with an *e*-value > 0.05 or less than 10 associated sequences were discarded and the sequences associated with the top five enriched motifs were used to construct PWMs. The “signif” PWM set was defined as the PWM for each TF with the most significant *e*-value, whereas in the “best” PWM set the top five most significant PWM was used for scoring and the PWM that gave off the highest variant effect prediction was used. All PWMs were constructed using the `toPWM` function of the `TFBStools` package^51^ and the `PWMscoreStartingAt` function of the `Biostrings` package was used to score sequences using the generated PWMs^52^.

The gkmtrain command from the LS-GKM library (https://github.com/Dongwon-Lee/lsgkm) was used to train gkmSVM models^12^ with default parameters, except for word length option set to 10. ChIP-seq and SELEX sequences were used as positive sequences, and random genomic sequences with the same length were used as negative sequences. The *deltaSVM* scores were generated from using the `gkmpredict` command along with the `deltasvm.pl` script (http://www.beerlab.org/deltasvm/). Finally, pre-trained GERV models for a total of 60 TFs were obtained from http://gerv.csail.mit.edu/ ChIP-seq experiments from ENCODE project and the preprocess and score options of the run.r script with default options was used to score the impact of variants.

### 1.6 Variant impact scoring metrics

#### 1.6.1 DeepBind and PWMs

Given a PWM or DeepBind model, we define a fixed-length sequence window of size *k* using the width of the PWM or the detector length of the DeepBind model (see^5^), respectively. Given a variant at position *q*, we score both the wildtype and mutant sequences starting *q* – *k* to *q* + *k* at increments of *k* for a set of raw wildtype scores *w*_1_, *w*_2_,…, *w_k_* and mutant scores *m*_1_,…, *m_k_*. Given the set of indices *S* = 1,…, *k*, the *delta raw* (Δ*R*) and *delta track* (Δ*T*) metrics are computed as follows:

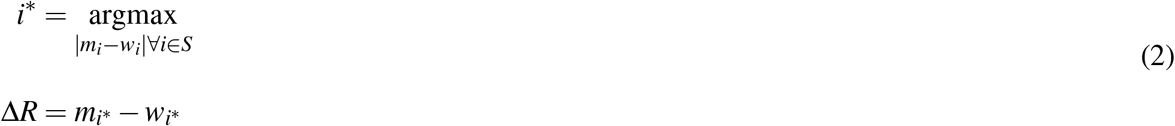

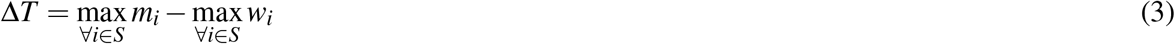

To compute *P_bind_* scores we score a given an individual raw score, we compute a foreground and background distribution of raw scores using a set of positive and negative sequence respectively. The negative sequences are defined as 10,000 randomly sampled genomic sequences of size k. The positive sequences for JASPAR PWMs are defined as generated sequences from the PWM, whereas for MEME-ChIP PWMs this is defined as the corresponding matching ChIP-seq sequences used to construct the PWM. The positive sequences as the ChIP-seq or SELEX sequences used to train the DeepBind models. We assume the background distribution follows a Gaussian distribution *N* ∼ *N* (*μ_n_*, σ_*n*_) and learn the parameters of the true positive distributions by fitting a two-component Gaussian model mixture model.

Here, one component was fixed to *μ_n_*, *σ_n_* and the true positive parameters are learned as *μ_p_*, *σ_p_*. A total of 10,000 random samples are generated using the given the parameters of background and used to train a generalised linear model, which was used to compute a posterior probability (**Supplementary Figure S3**) of binding (*P_bind_*) and not binding (*P_not binding_*). The *P_bind_* scores are computed for both the wildtype 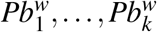 and mutant 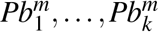 windows. The *P_not binding_* scores are computed similarly as 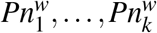 and 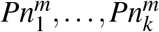. The delta *P_bind_* (Δ*P*) score was then computed similarly to that of Δ*R*:

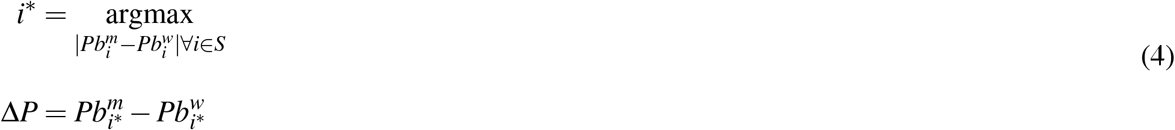

The probabilistic scores of a loss (*P_loss_*) and gain (*P_gain_*) events are computed by multiplying the likelihood of the wildtype allele binding and the mutant allele not binding for loss, and vice versa for gain:

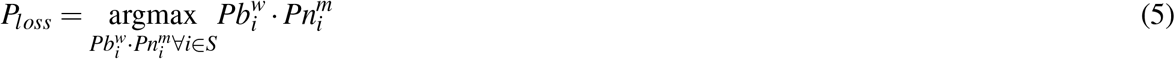

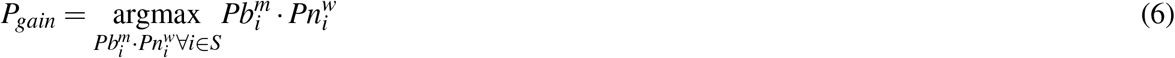

Both probabilities are then combined into individual scores *P_sum_* and *P_comb_* as follows:

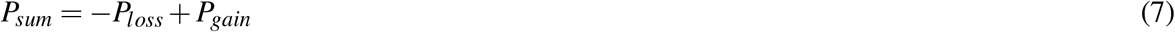

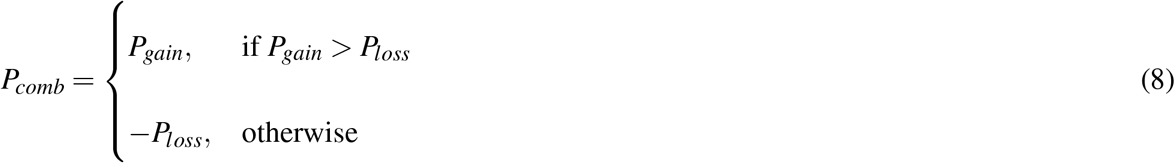

#### 1.6.2 DeepSEA

Given a probability of binding in the wildtype and mutant alleles, as *ρ_w_* and *ρ_m_* respectively, DeepSEA utilises two scoring schemes: the difference (*DS_D_*) and log fold change (*DS_L_*) computed as follows:

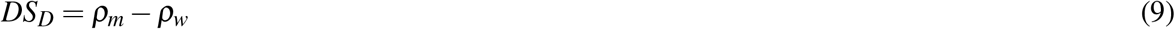

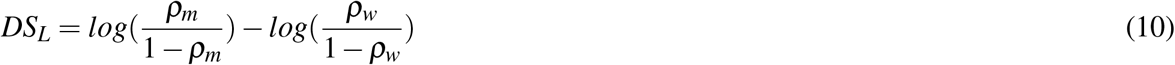

#### 1.6.3 gkmSVM

Given *k*-mer weights computed for wildtype and mutant sequences flanking the variant position as 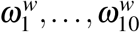 and 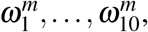 the deltaSVM (Δ*SVM*) score is computed as follows:

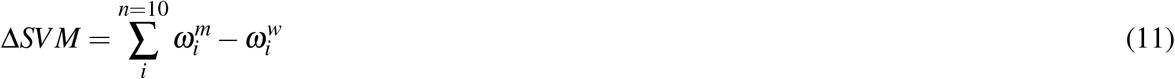

### 1.7 Allele frequencies and non-coding variant impact predictions

Allele frequencies for the 1000 genomes project, ExAC variants and ESP6500 project, along with non-coding variant impact predictions for CADD, Eigen and GWAVA were obtained from the ANNOVAR tool^53^, using the `table_annovar.pl` script.

### 1.8 Performance measures

ROC and PR curves were generated by assessing the TPR (or recall), FPR and precision. The ROC curves compare the FPR against the TPR, whereas PR curves compare the TPR against the PPV (or precision). Given the number of true positives (TP), false positives (FP), true negatives (TN) and false negatives (FN), these are computed as follows:

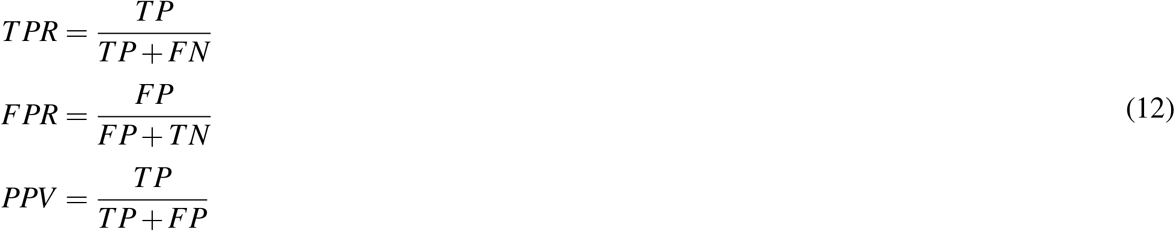

All ROC and PR curves, along with area under the curve measures were computed using the PRROC R package^54^.

## Author contributions statement

O.W., D.M. and A.D. conceived the study, O.W. conducted and analysed the results. All authors reviewed the manuscript.

## References

1. Sandelin, A., Alkema, W., Engström, P., Wasserman, W. W. & Lenhard, B. JASPAR: an open-access database for eukaryotic transcription factor binding profiles. Nucleic acids research 32, D91–4 (2004). DOI 10.1093/nar/gkh012.

2. Tehranchi, A. K. et al. Pooled ChIP-Seq Links Variation in Transcription Factor Binding to Complex Disease Risk. Cell 165,730–41 (2016). DOI 10.1016/j.cell.2016.03.041.

3. Kilpinen, H. et al. Coordinated effects of sequence variation on DNA binding, chromatin structure, and transcription. Sci. (New York, N.Y.) 342, 744–7 (2013). DOI 10.1126/science.1242463.

4. Wasserman, W. W. & Sandelin, A. Applied bioinformatics for the identification of regulatory elements. Nat. reviews. Genet. 5, 276–87 (2004). DOI 10.1038/nrg1315.

5. Alipanahi, B., Delong, A., Weirauch, M. T. & Frey, B. J. Predicting the sequence specificities of DNA- and RNA-binding proteins by deep learning. Nat. biotechnology 33, 831–8 (2015). DOI 10.1038/nbt.3300.

6. Zhou, J. & Troyanskaya, O. G. Predicting effects of noncoding variants with deep learning-based sequence model. Nat. methods 12, 931–4 (2015). DOI 10.1038/nmeth.3547.

7. McLaren, W. et al. The Ensembl Variant Effect Predictor. Genome biology 17, 122 (2016). DOI 10.1186/s13059-016-0974-4.

8. Weirauch, M. T. et al. Evaluation of methods for modeling transcription factor sequence specificity. Nat. biotechnology 31, 126–34 (2013). DOI 10.1038/nbt.2486.

9. Jayaram, N., Usvyat, D. & R Martin, A. C. Evaluating tools for transcription factor binding site prediction. BMC bioinformatics (2016). DOI 10.1186/s12859-016-1298-9.

10. Kaplun, A. et al. Establishing and validating regulatory regions for variant annotation and expression analysis. BMC genomics 17 Suppl 2, 393 (2016). DOI 10.1186/s12864-016-2724-0.

11. Castro, M. A. A. et al. Regulators of genetic risk of breast cancer identified by integrative network analysis. Nat. genetics 48, 12–21 (2016). DOI 10.1038/ng.3458.

12. Lee, D. et al. A method to predict the impact of regulatory variants from DNA sequence. Nat. genetics 47, 955–61 (2015). DOI 10.1038/ng.3331.

13. Zeng, H., Hashimoto, T., Kang, D. D. & Gifford, D. K. GERV: a statistical method for generative evaluation of regulatory variants for transcription factor binding. Bioinforma. (Oxford, England) 32, 490–6 (2016). DOI 10.1093/bioinformatics/btv565.

14. Shi, W., Fornes, O., Mathelier, A. & Wasserman, W. W. Evaluating the impact of single nucleotide variants on transcription factor binding. Nucleic acids research 44, 10106–10116 (2016). DOI 10.1093/nar/gkw691.

15. de Santiago, I. et al. BaalChIP: Bayesian analysis of allele-specific transcription factor binding in cancer genomes. Genome biology 18, 39 (2017). DOI 10.1186/s13059-017-1165-7.

16. Chen, J. et al. A uniform survey of allele-specific binding and expression over 1000-Genomes-Project individuals. Nat. communications 7, 11101 (2016). DOI10.1038/ncomms11101.

17. Reddy, T. E. et al. Effects of sequence variation on differential allelic transcription factor occupancy and gene expression. Genome research 22, 860–9 (2012). DOI 10.1101/gr.131201.111.

18. Wei, Y., Li, X., Wang, Q.-f. & Ji, H. iASeq: integrative analysis of allele-specificity of protein-DNA interactions in multiple ChIP-seq datasets. BMC genomics 13, 681 (2012). DOI 10.1186/1471-2164-13681.

19. Bailey, S. D., Virtanen, C., Haibe-Kains, B. & Lupien, M. ABC: a tool to identify SNVs causing allele-specific transcription factor binding from ChIP-Seq experiments. Bioinforma. (Oxford, England) 31, 3057–9 (2015). DOI 10.1093/bioinformatics/btv321.

20. Lek, M. et al. Analysis of protein-coding genetic variation in 60,706 humans. Nat. 536, 285–91 (2016). DOI 10.1038/nature19057.

21. Siva, N. 1000 Genomes project. Nat. biotechnology 26, 256 (2008). DOI 10.1038/nbt0308-256b.

22. Fu, W. et al. Analysis of 6,515 exomes reveals the recent origin of most human protein-coding variants. Nat. 493, 216–20 (2013). DOI 10.1038/nature11690.

23. Yuen, R. K. C. et al. Genome-wide characteristics of de novo mutations in autism. NPJ genomic medicine 1, 160271–1602710 (2016). DOI10.1038/npjgenmed.2016.27.

24. Ritchie, G. R. S., Dunham, I., Zeggini, E. & Flicek, P. Functional annotation of noncoding sequence variants. Nat. methods 11, 294–6 (2014). DOI 10.1038/nmeth.2832.

25. Ionita-Laza, I., McCallum, K., Xu, B. & Buxbaum, J. D. A spectral approach integrating functional genomic annotations for coding and noncoding variants. Nat. genetics 48, 214–20 (2016). DOI 10.1038/ng.3477.

26. Kircher, M. et al. A general framework for estimating the relative pathogenicity of human genetic variants. Nat. genetics 46, 310–5 (2014). DOI 10.1038/ng.2892.

27. Khurana, E. et al. Integrative annotation of variants from 1092 humans: application to cancer genomics. Sci. (New York, N.Y.) 342, 1235587 (2013). DOI 10.1126/science.1235587.

28. Fu, Y. et al. FunSeq2: a framework for prioritizing noncoding regulatory variants in cancer. Genome biology 15, 480 (2014). DOI 10.1186/s13059-014-0480-5.

29. Perera, D. et al. OncoCis: annotation of cis-regulatory mutations in cancer. Genome biology 15, 485 (2014). DOI 10.1186/s13059-014-0485-0.

30. Payne, J. L. & Wagner, A. Mechanisms of mutational robustness in transcriptional regulation. Front. genetics 6, 322 (2015). DOI 10.3389/fgene.2015.00322.

31. Machanick, P. & Bailey, T. L. MEME-ChIP: motif analysis of large DNA datasets. Bioinforma. (Oxford, England) 27, 1696–7 (2011). DOI 10.1093/bioinformatics/btr189.

32. Melton, C., Reuter, J. A., Spacek, D. V. & Snyder, M. Recurrent somatic mutations in regulatory regions of human cancer genomes. Nat. genetics 47, 710–6 (2015). DOI 10.1038/ng.3332.

33. Slattery, M. et al. Cofactor binding evokes latent differences in DNA binding specificity between Hox proteins. Cell 147, 1270–82 (2011). DOI 10.1016/j.cell.2011.10.053.

34. Schaefer, U., Schmeier, S. & Bajic, V. B. TcoF-DB: dragon database for human transcription co-factors and transcription factor interacting proteins. Nucleic acids research 39, D106–10 (2011). DOI 10.1093/nar/gkq945.

35. Yin, Y. et al. Impact of cytosine methylation on DNA binding specificities of human transcription factors. Sci. (New York, N.Y.) 356 (2017). DOI 10.1126/science.aaj2239.

36. Spruijt, C. G. et al. Dynamic readers for 5-(hydroxy)methylcytosine and its oxidized derivatives. Cell 152, 1146–59 (2013). DOI 10.1016/j.cell.2013.02.004.

37. Zelko, I. N., Mueller, M. R. & Folz, R. J. CpG methylation attenuates Sp1 and Sp3 binding to the human extracellular superoxide dismutase promoter and regulates its cell-specific expression. Free. radical biology & medicine 48, 895–904 (2010). DOI 10.1016/j.freeradbiomed.2010.01.007.

38. Rohs, R. et al. The role of DNA shape in protein-DNA recognition. Nat. 461, 1248–53 (2009). DOI 10.1038/nature08473.

39. Mathelier, A. et al. DNA Shape Features Improve Transcription Factor Binding Site Predictions In Vivo. Cell systems 3, 278–286.e4 (2016). DOI 10.1016/j.cels.2016.07.001.

40. Whitmarsh, A. J. & Davis, R. J. Regulation of transcription factor function by phosphorylation. Cell. molecular life sciences: CMLS 57, 1172–83 (2000).

41. Hornbeck, P. V. et al. PhosphoSitePlus, 2014: mutations, PTMs and recalibrations. Nucleic acids research 43, D512–20 (2015). DOI 10.1093/nar/gku1267.

42. Stringham, J. L., Brown, A. S., Drewell, R. A. & Dresch, J. M. Flanking sequence context-dependent transcription factor binding in early Drosophila development. BMC bioinformatics 14, 298 (2013). DOI 10.1186/1471-2105-14-298.

43. Gordân, R. et al. Genomic regions flanking E-box binding sites influence DNA binding specificity of bHLH transcription factors through DNA shape. Cell reports 3, 1093–104 (2013). DOI 10.1016/j.celrep.2013.03.014.

44. Domcke, S. et al. Competition between DNA methylation and transcription factors determines binding of NRF1. Nat. 528, 575–9 (2015). DOI 10.1038/nature16462.

45. Pique-Regi, R. et al. Accurate inference of transcription factor binding from DNA sequence and chromatin accessibility data. Genome research 21, 447–55 (2011). DOI 10.1101/gr.112623.110.

46. Waterman, M. J., Stavridi, E. S., Waterman, J. L. & Halazonetis, T. D. ATM-dependent activation of p53 involves dephosphorylation and association with 14-3-3 proteins. Nat. genetics 19, 175–8 (1998). DOI 10.1038/542.

47. Arvey, A., Agius, P., Noble, W. S. & Leslie, C. Sequence and chromatin determinants of cell-type-specific transcription factor binding. Genome research 22, 1723–34 (2012). DOI 10.1101/gr.127712.111.

48. Vaquerizas, J. M., Kummerfeld, S. K., Teichmann, S. A. & Luscombe, N. M. A census of human transcription factors: function, expression and evolution. Nat. reviews. Genet. 10, 252–63 (2009). DOI 10.1038/nrg2538.

49. Kulakovskiy, I. V. et al. HOCOMOCO: a comprehensive collection of human transcription factor binding sites models. Nucleic acids research 41, D195–202 (2013). DOI10.1093/nar/gks1089.

50. Kent, W. J. et al. The human genome browser at UCSC. Genome research 12, 996–1006 (2002). DOI10.1101/gr.229102. Article published online before print in May 2002.

51. Tan, G. & Lenhard, B. TFBSTools: an R/bioconductor package for transcription factor binding site analysis. Bioinforma. (Oxford, England) 32, 1555–6 (2016). DOI 10.1093/bioinformatics/btw024.

52. Pagès, H., Aboyoun, P., Gentleman, R. & DebRoy, S. Biostrings: String objects representing biological sequences, and matching algorithms (2016). R package version 2.42.1.

53. Wang, K., Li, M. & Hakonarson, H. ANNOVAR: functional annotation of genetic variants from high-throughput sequencing data. Nucleic acids research 38, e164 (2010). DOI 10.1093/nar/gkq603.

54. Grau, J., Grosse, I. & Keilwagen, J. PRROC: computing and visualizing precision-recall and receiver operating characteristic curves in R. Bioinforma. (Oxford, England) 31, 2595–7 (2015). DOI 10.1093/bioinformatics/btv153.

